# Maternal separation stress experienced by parents affects ovarian function in first generation of mice

**DOI:** 10.1101/2020.02.26.966192

**Authors:** Kajal Khodamoradi, Zahra Khosravizadeh, Hossein Amini-Khoei, Seyed Reza Hosseini, Ahmad Reza Dehpour, Gholamreza Hassanzadeh

**Author notes:** Correspondence: Kajal Khodamoradi: Department of Anatomy, School of Medicine, Tehran University of Medical Sciences, Tehran, Iran, And Emai.

## Abstract

The maternal separation stress during postnatal development can adversely affect one’s adulthood. Some parents’ experiences may not only affect the phenotype of parents but also alter the reaction to environmental impacts in the offspring. The aim of this study is to investigate consequences of maternal separation stress in female first generation of mice whose parents were exposed to maternal separation stress. Maternal separation in pups was performed during post-natal days (PND) 2 to 14. Then, female pups of the first-generation were used in present study. The histological changes in ovaries, ROS production (using DCFH-DA assay), mRNA expression of NLRP3, ASC, caspase-1, TLR4, BAX, BCL2 and TNFα genes (using RT-PCR), levels of IL-18, IL-1β, ATP and GPx (using ELISA) and also protein expression of caspase-3 and NLRP3 (using immunocytochemistry) were assessed. Our findings showed that maternal separation stress experienced by parents significantly affects the numbers of primordial and primary follicles. Furthermore, ROS production increased and concentrations of ATP and GPx reduced in the first generation. Also, expression of cytokines and genes involved in inflammation and apoptosis including NLRP3, caspase-1, TLR4, TNFα, IL-1β, IL-18 and BCL2 were significantly affected in the first generation. Our results also showed that this stress significantly increased percentage of caspase-3 and NLRP3 positive cells in the ovarian tissue of the first generation. Our findings suggest that maternal separation stress experienced by parents may influence activation of inflammatory response in the ovarian tissue of their first generation which may induce apoptosis and consequently disturb folliculogenesis process.

## Introduction

It is well known that the mother-offspring relationship during the neonatal period is required for proper development because mother acts as the communication loop between the first environment and development of animal (1). The maternal separation stress during postnatal development can have adversely effects throughout adulthood (2). Furthermore, the results of studies indicated that early-life stress can play a role in causing a variety of psychiatric disorders in the future (3). Indeed, stress can disturb the function of gonad not only in humans but also in all higher animals. Stress-induced gonadal dysfunction contains impairment in the function of hypothalamic-pituitary-gonadal axis (4). It is appeared that repeated activation of the hypothalamic-pituitary adrenal (HPA) axis can lead to disharmony in the neurotrophic and immune pathways. This disharmony can have a main role in the long-term effects of early-life stress (5). In addition, stress can induce the activation of the adrenergic system that alter the function of the immune and endocrine systems. Cooperation between these two systems is essential for normal female fertility (4, 6).

Several studies suggested that different factors including danger signals, catecholamines and glucocorticoids play important roles in stimulation of immune system during stress (7-9). Inflammasomes are intracellular protein complexes which play main roles in the innate immune and reproductive systems in mammals (10). NLRP3 inflammasome activation, as a well-known NLR, results in the secretion of the pro-inflammatory cytokines IL-1β and IL-18 (11). It is demonstrated that maternal separation stress can activate the NLRP3 inflammasome components and consequently IL-1β and IL-18 (12). In previous study, we studied the effect of stress-induced maternal separation on the ovarian function in adult female mice. Our findings showed that maternal separation stress has adverse effects on ovarian tissue. Harmful effects are probably occurring through increase of ROS production and impact on mitochondrial function, inflammatory process and apoptosis pathways.

According to epidemiological investigations, the offspring of persons with stress-induced changes in behavior, and sometimes their next generation, can be also affected even if they did not directly experience the same stressful event (13-16). It is evidenced that some parents’ experiences may not only affect the phenotype of parents but also alter the reaction to environmental impacts in the offspring (14). To our knowledge, the effect of maternal separation on reproductive potential across generations has not been deeply evaluated. Therefore, this study was designed to investigate consequences of maternal separation stress in female first generation of mice whose parents were exposed to maternal separation stress.

## Materials and methods

### Experimental animals

Pregnant NMRI (Naval Medical Research Institute) mice were obtained from the Pasteur Institute of Iran. The animals were housed under controlled temperature (22 - 25°C) and humidity (55 - 65%), with a light/dark cycle of 12 h: 12 h. All procedures were carried out according to guidelines approved by the Ethics Committee of Tehran University of Medical Sciences (IR.TUMS.MEDICINE.REC.1395.2507). In this experimental study, pups were randomly allocated to two groups: maternal separation and control group. In maternal separation group, pups were separated from their mothers and placed in a separate, clean cage for 3 hours every day (9 am to 12 pm each day) from postnatal day (PND) 2 to 14. The day of birth was considered as PND 0 (17-19). The pups of the control group were left untouched. Pups of the first-generation litters. Then, female pups of first-generation were used in present study. Mice at PND 70 were sacrificed under deep anesthesia and the ovaries were removed. One ovary from each mouse was used for molecular assessments and the second ovary was used for histological assessment. All experiments were performed in triplicate and repeated three times.

### Histological evaluation

The ovarian tissues for histological evaluation were directly transferred to Bouin’s fixative solution. The fixed ovaries were dehydrated through ascending graded series of ethanol (Merck, Darmstadt, Germany) and then placed in paraffin wax. The serial sections, 5 μm thick, were prepared using a rotary microtome (Microm, Walldorf, Germany) and rehydrated through descending graded series of ethanol. The sections were dewaxed in xylene, stained with haematoxylin and eosin and mounted with DPX. For histological evaluation, transverse sections from nine different regions of the ovaries were examined. The number of primordial, primary, secondary and graafian follicles were counted by light microscopy and ImageJ software (ImageJ U. S. National Institutes of Health, Bethesda, MD, USA).

### ROS assay

The level of ROS production in ovarian tissues was measured with flow cytometry using 2′, 7′ -dichlorofluorescin diacetate (DCFH-DA; Sigma, USA) after enzymatic digestion of minced tissue (20). The ovarian tissues were mechanically homogenized in Ham’s F-10 medium (Life Technology, Carlsbad, CA, USA). Tissue homogenates were centrifuged at 10,000 x g for 5 min and washed with PBS. After incubation of the homogenates with 20 µM DCFH-DA in dark at 37°C for 45 min, tissue homogenates were washed with PBS. Then, DCF fluorescence (green) was detected in the FL-1 channel using a BD FACScan flow cytometer (Becton Dickinson, San Jose, CA, USA)(21).

### Real-time reverse transcription polymerase chain reaction (RT-qPCR) analysis

The level of mRNA expressions of NLRP3, ASC, caspase-1, TLR4, BAX, BCL2 and TNFα genes was analyzed by RT-qPCR. The total RNA extraction from ovarian tissues was performed with TRIzol reagent according to the manufacturer’s protocol (Invitrogen, Carlsbad, CA). The complementary DNA (cDNA) was synthesis via reverse transcription reaction using a PrimeScript RT reagent kit (Takara, South Korea) according to the manufacturer’s instructions. The RT-qPCR was carried out with gene specific primers and the HOT FIREPol EvaGreen qPCR Mix Plus (Solis BioDyne, Tartu, Estonia) by an ABI7500 (Applied Biosystems, Foster City, California, USA). The normalization of mRNA expression levels was performed using the reference gene glyceraldehydes-3-phosphate dehydrogenase (GAPDH) mRNA expression. The mRNA levels of target genes expression was calculated using 2^-ΔCT^. List of primer sequences used for RT-qPCR analysis are listed in Table 1.

**Table 1.**
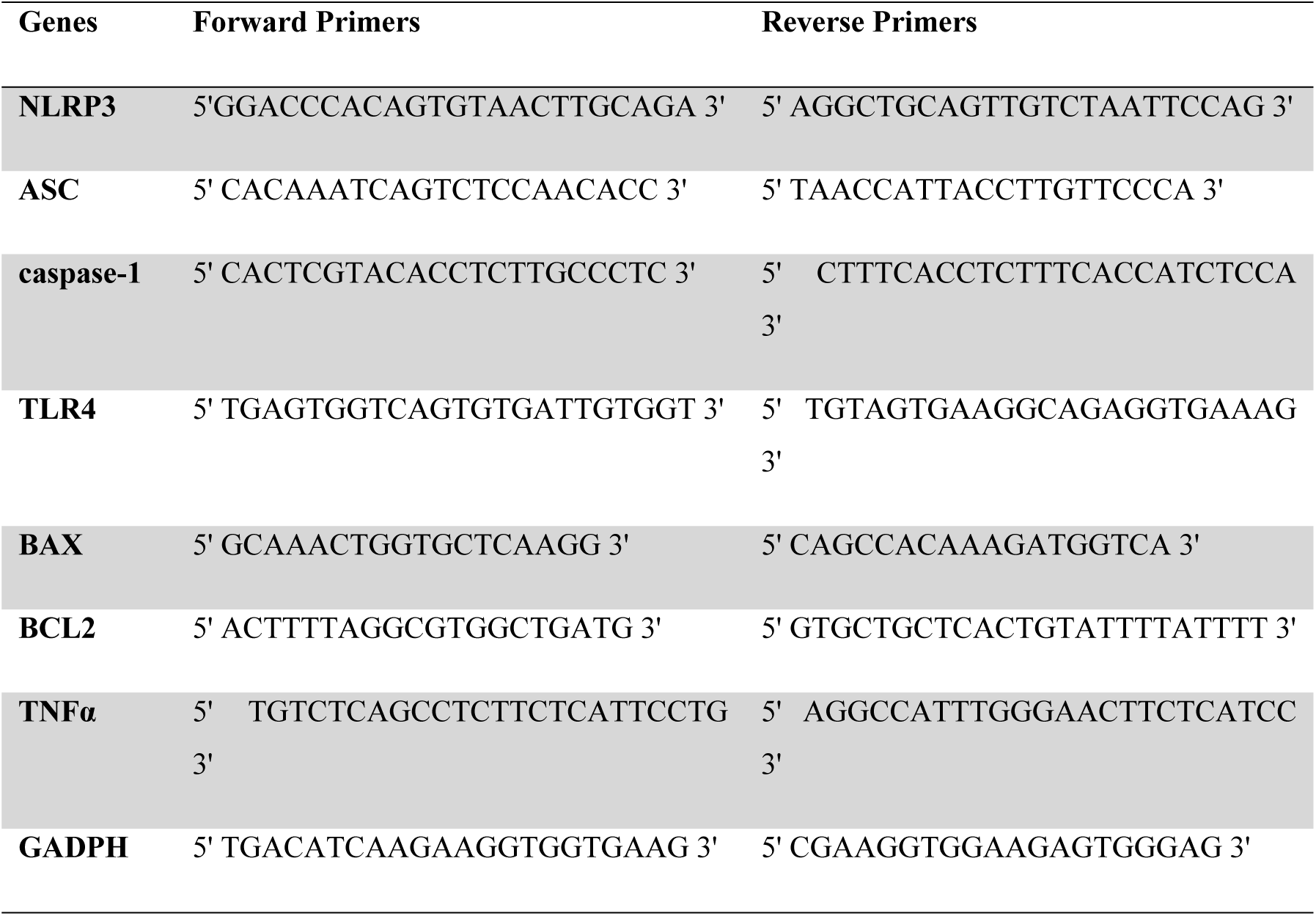
Primer Sequences.

### Enzyme-linked immunosorbent assay (ELISA)

The specimen of ovarian tissues were homogenized in PBS on ice, then centrifuged at 800 × g for 5 min. The harvested supernatants were used in the ELISA assay. The level concentrations of IL-18 and IL-1β in ovarian tissue were measured using ELISA kits (Koma Biotech–Korea) and the level concentrations of ATP and GPx were measured with specific ELISA kits (R&D Systems, Minneapolis, MN, USA) according to the manufacturer’s protocols.

### Immunocytochemical analysis

To assess caspase-3 and NLRP3 immunoreactivity, the tissue specimens were dehydrated in a series of graded ethanol and embedded in paraffin. After removing paraffin, tissue specimens were rehydrated in graded series of ethanol and permeabilized with 10 mM sodium citrate and 0.05% Tween 20. The samples were blocked in a blocking solution including 1% (w/v) bovine serum albumin (BSA; Sigma-Aldrich, St. Louis, MO, USA) in PBS. Then, the ovarian samples were incubated overnight at 4°C with primary antibodies against caspase-3 (1:1000 dilution, Abcam, Cambridge, MA) and NLRP3 (1:1000 dilution, Abcam, Cambridge, MA). After incubation of the samples with secondary antibody (1:500 dilution, Abcam, Cambridge, MA) for 2 hour at 37°C, the cells’ nuclei were stained with PI (1:1000, Sigma-Aldrich, St. Louis, MO, USA). The cell counting and merging of the pictures were performed by “Image J” software (Image J U. S. National Institutes of Health, Bethesda, Maryland, USA).

### Statistical Analysis

The collected data were analyzed using SPSS version 20.0 software. The normality of the values were tested using the Kolmogorov–Smirnov test. The independent samples t-test and Mann-Whitney U test were used to determine the statistical significance of the results. The results were presented as the mean ± SD (standard deviation) and p ≤ 0.05 was considered statistically significant.

## Results

### Histological evaluation

Histological analysis of ovarian follicles was evaluated with haematoxylin and eosin (H & E) staining (Fig. 1). The results showed that percentage of primordial follicles in the first-generation group (20.60 ± 12.0332) was significantly lower compared with the control group (50.5812 ± 1.6164) (p < 0. 05). There was significant difference between percentage of primary follicles in the first-generation group (14.2182 ± 4.0577) and percentage of primary follicles in the control group (3.4095 ± 0.0548) (p < 0. 05). There was no significant difference between percentage of secondary follicles in the first-generation group (24.4364 ± 4.4412) and percentage of secondary follicles in the control group (22.7173 ± 1.2419, p > 0. 05). In addition, percentage of graafian follicles in the first-generation group (40.7455± 18.3935) was higher compared with the control group (23.2920 ± 0.4292, p > 0. 05, Fig.2), although this difference was not significant.

**Fig. 1.**
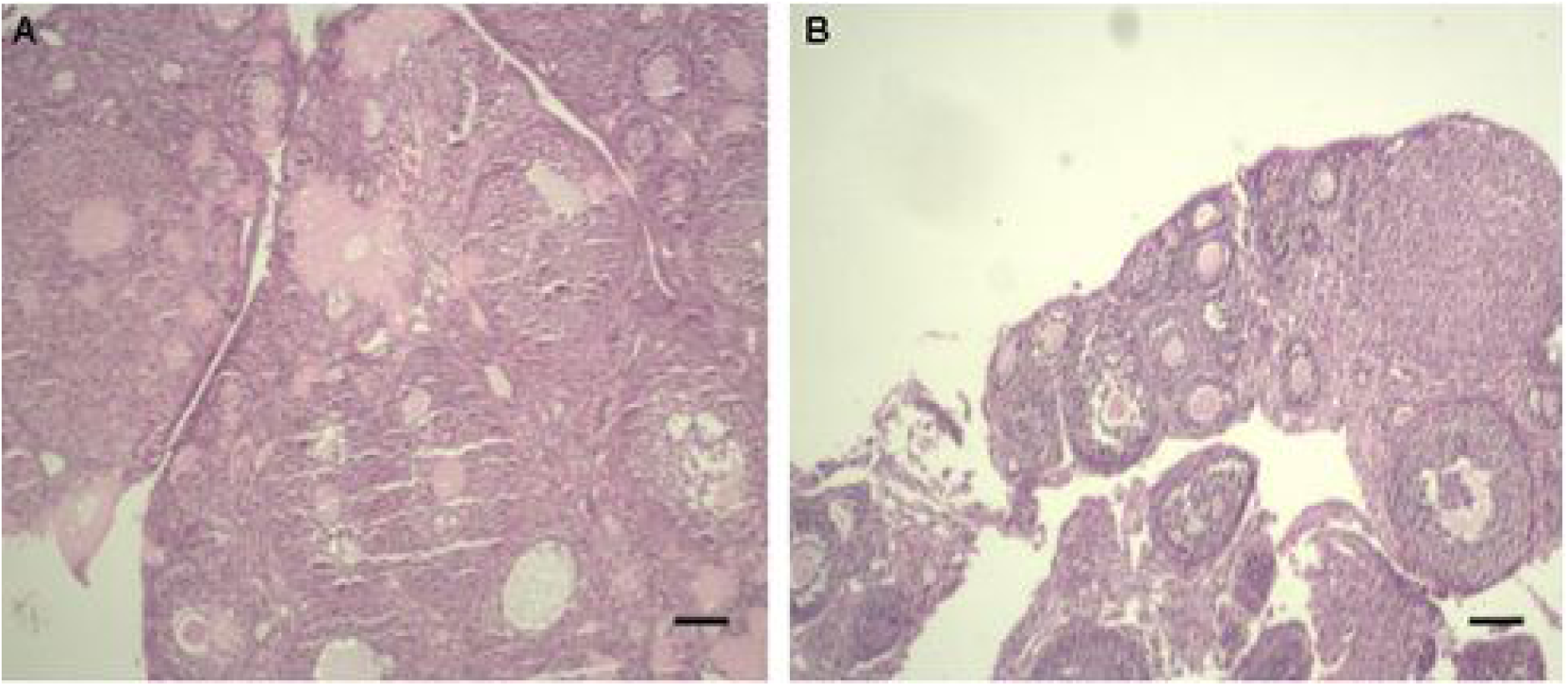
The histopathological features provided from H & E-stained ovarian sections in the control (A) and the first-generation (B) groups. Scale bars are 10 µm.

**Fig.2.**
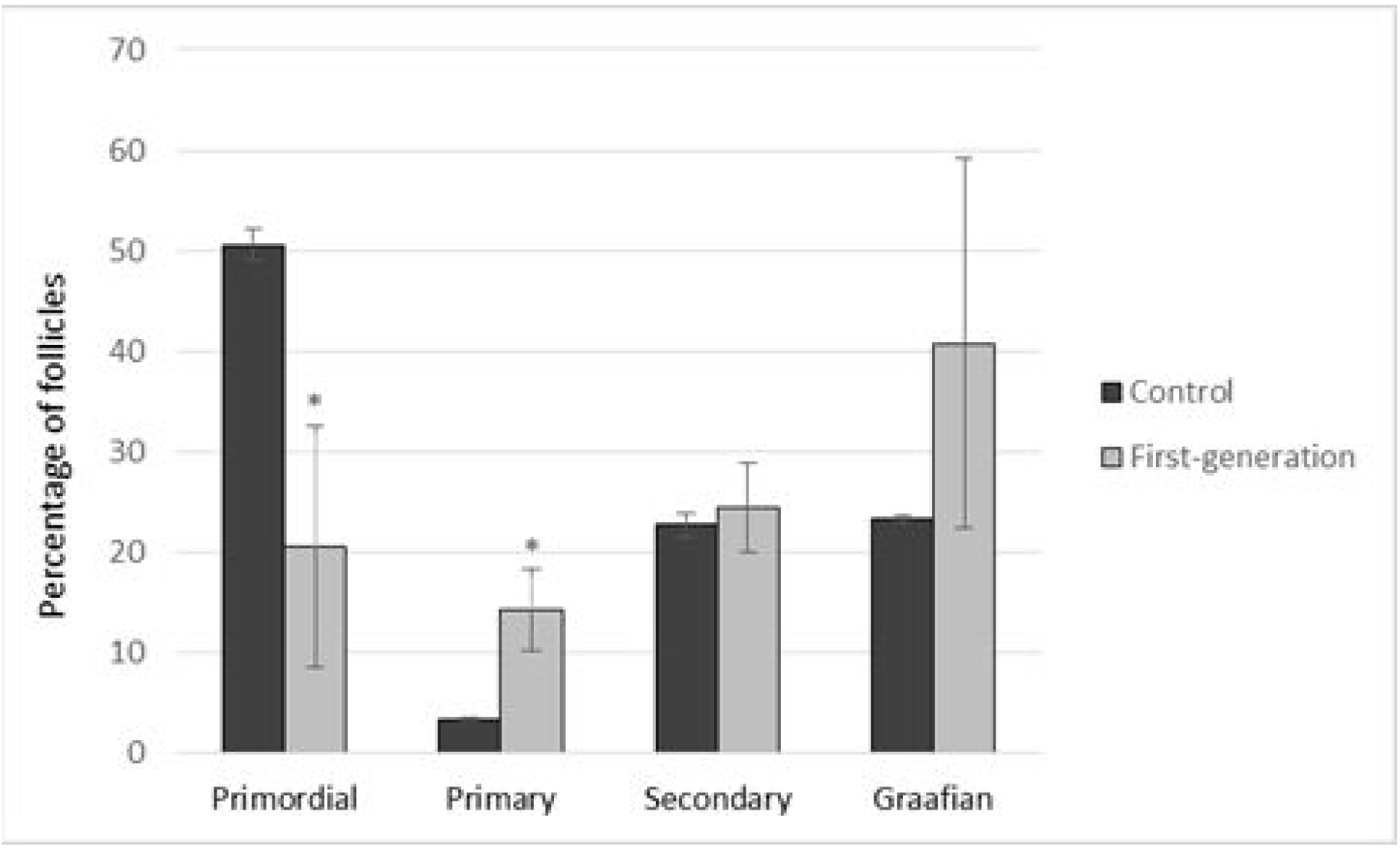
Effect of maternal separation on mice ovarian follicles in the first-generation. Values are reported as mean ± SD. *p < 0.05 compared with the control group.

### ROS evaluation

ROS production in ovarian tissue was detected using the DCFH-DA assay. The level of ROS production in ovarian tissue of the first-generation group (533.9831 ± 35.0721) was significantly higher compared with the control group (324. 8135 ± 36.4535, p < 0. 001, Fig. 3).

**Fig. 3.**
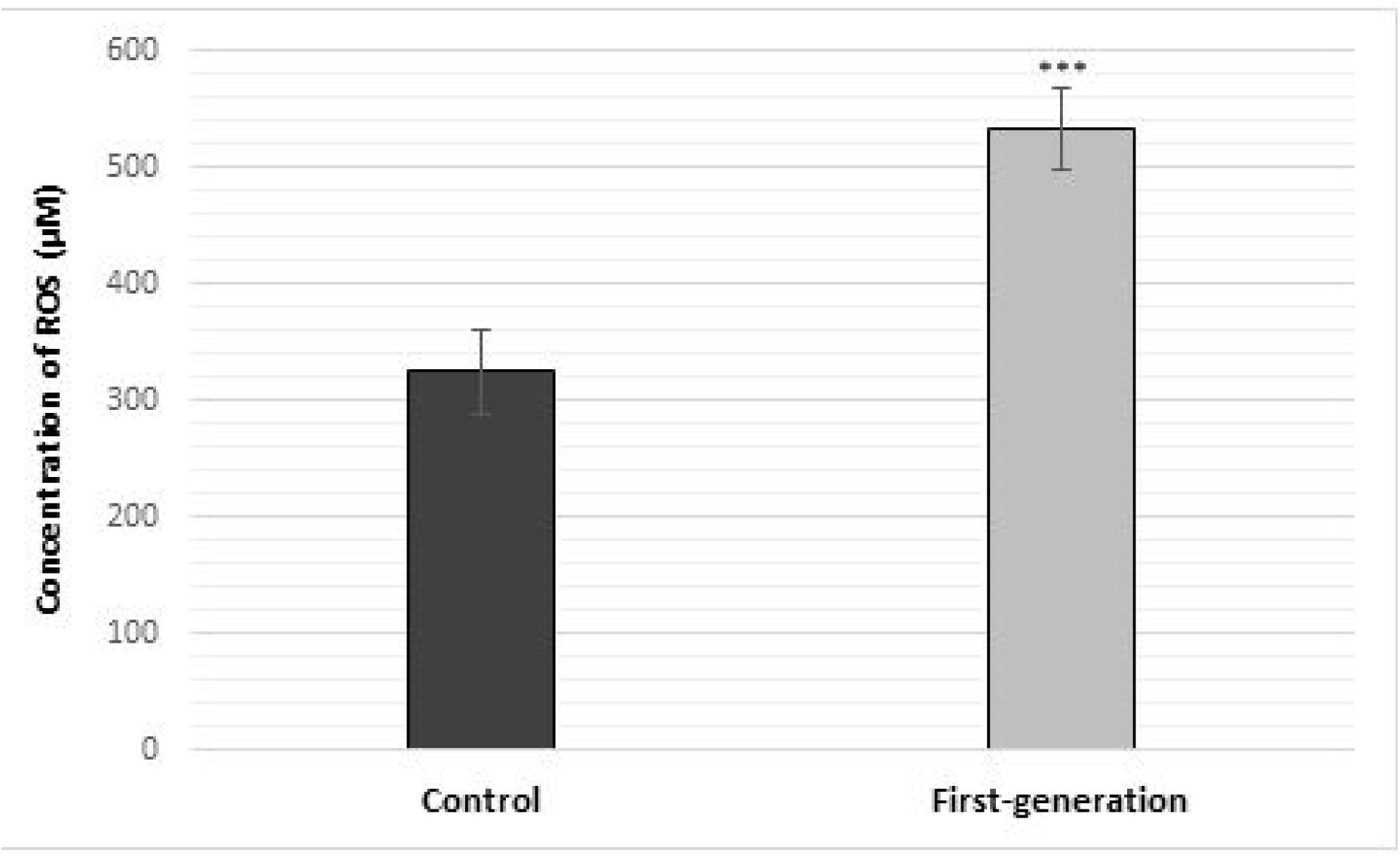
The level concentrations of ROS production in the ovary. Values are reported as mean ± SD. ***p < 0.001 compared with the control group.

### RT-qPCR analysis

The results of RT-qPCR analysis showed that expression of the NLRP3 and caspase-1 significantly increased in the first-generation group compared with the control group (p < 0.05). Also, expression of TLR4 and TNFα in ovarian tissue of the first-generation group was significantly higher compared with the control group (p < 0.01). In addition, expression of ASC and BAX in the first-generation group was higher compared with the control group, although this difference was not significant (p > 0. 05). In contrast, the expression of BCL2 gene significantly decreased in the first-generation group compared with the control group (p < 0.05) (Fig 4).

**Fig. 4.**
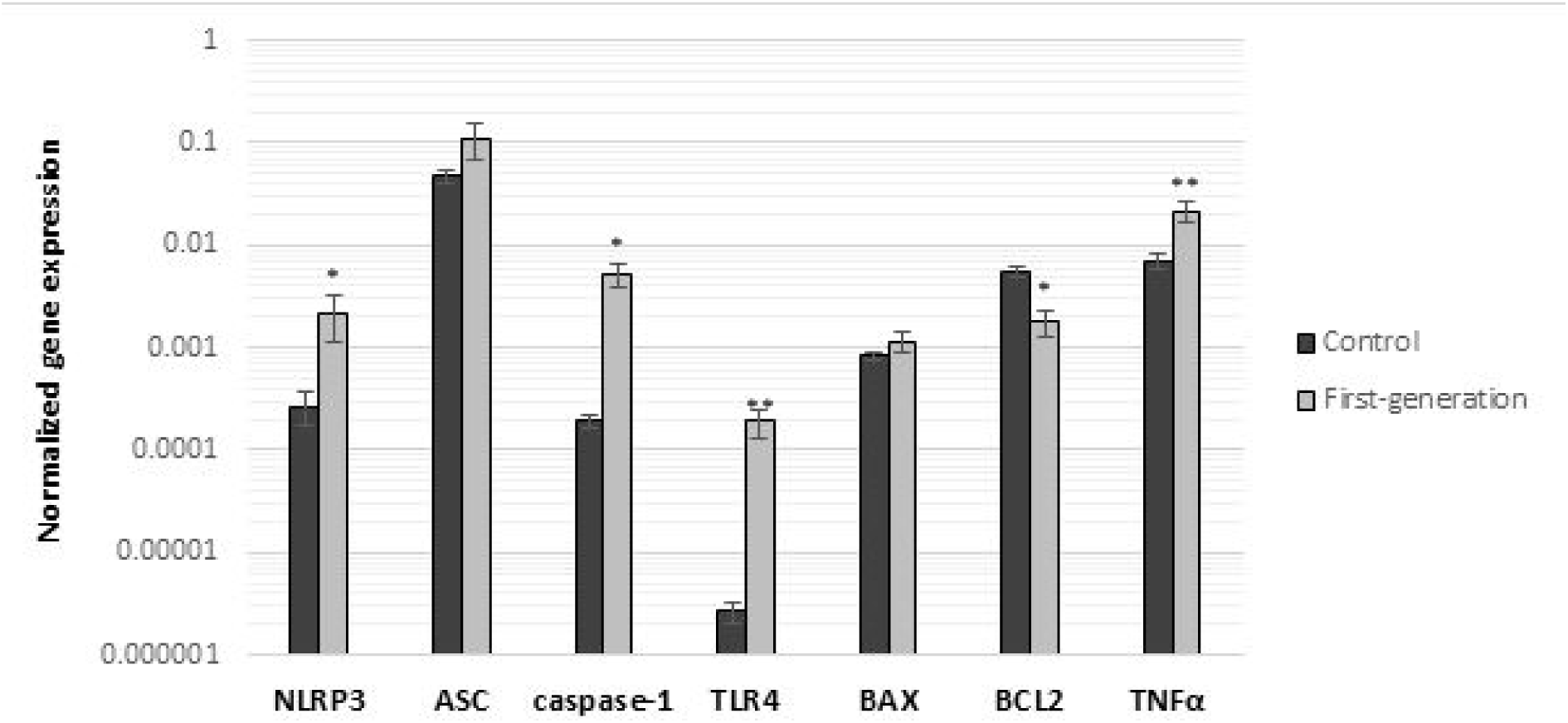
The gene expression of NLRP3, ASC, caspase-1, TLR4, BAX, BCL2 and TNFα using RT-qPCR. Values are reported as mean ± SD (Standard Deviation). *p < 0.0 and **p < 0.01 compared with the control group.

### Enzyme-linked immunosorbent assay (ELISA)

ELISA results showed that the level concentrations of IL-18 and IL-1β in the first-generation group were higher than the control group (2.6278 ± 0.06 vs. 2.1053 ± 0.07, p < 0.001 and 2.1328 ± 0.02 vs. 1.6285 ± 0.01, p < 0.001 respectively). The level concentrations of ATP and GPx were significantly lower in the first-generation group compared with the control group (2.6286 ± 0.04 vs. 2.8321 ± 0.03, and 212.9558 ± 14.23 vs. 304.0299 ± 15.6235 respectively, p < 0.01 for both) (Fig.5).

**Fig.5.**
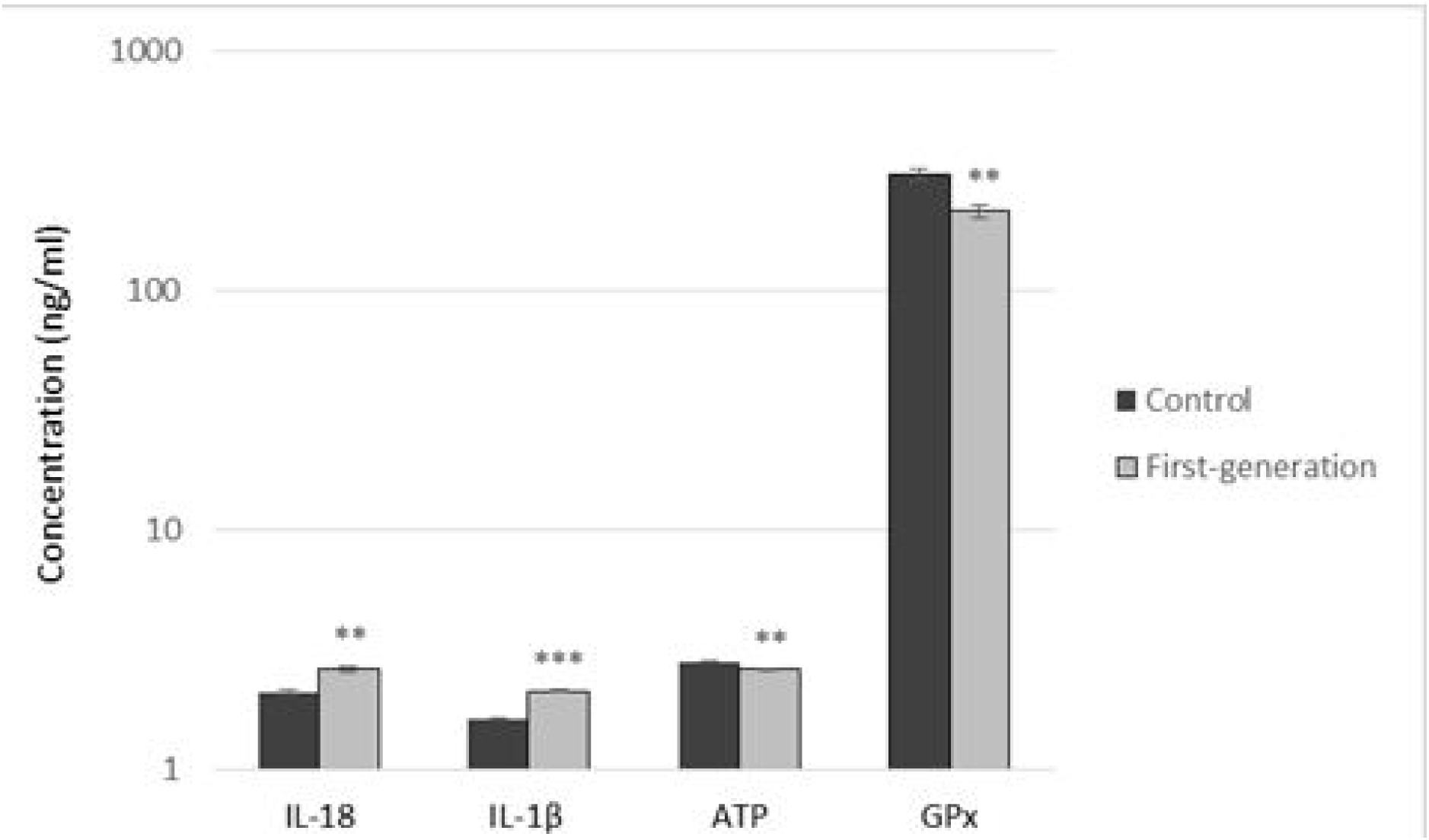
The level concentrations of IL-18, IL-1β, ATP and GPx measured using ELISA. Values are reported as mean ± SD. **p < 0.01 and ***p < 0.001 compared with the control group.

### Immunocytochemical analysis

Caspase-3 and NLRP3 markers were used to label the ovarian cells in the control and first-generation groups. The nuclei were stained with PI (Fig.6). Immunocytochemical analysis showed that the mean percentage of caspase-3 positive cells in the first-generation group was significantly higher compared to the control group (50.00 ± 1.00% vs. 29.50 ± 1.71%, p < 0.001). Also, the mean percentage of NLRP3 positive cells was significantly higher in the first-generation group in compared to the control group (47.00 ± 2.00 % vs. 27.00 ± 2.15 %, p < 0.001) (Fig.7).

**Fig 6.**
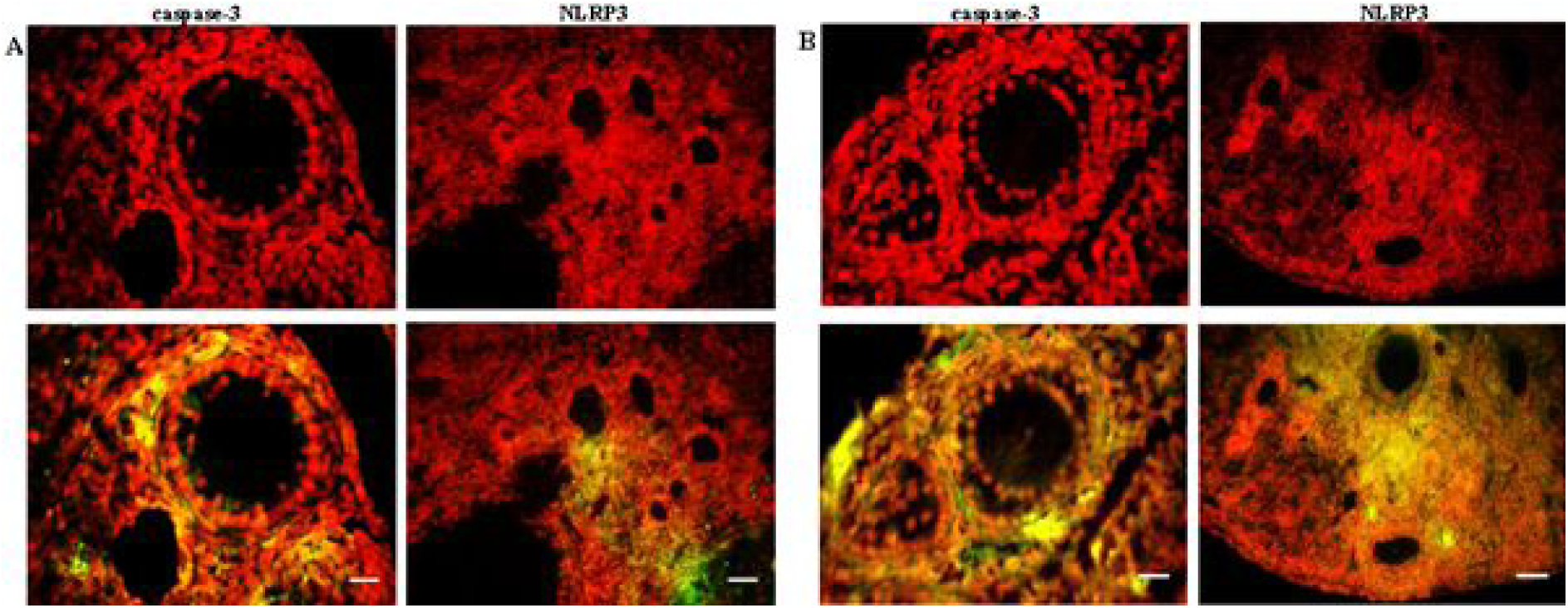
Immunocytochemical analysis of ovarian cells for caspase-3 and NLRP3 markers. (A) Control group; (B) first-generation group; upper panel: PI stained pictures; lower panel: merged pictures of PI and secondary antibody stained cells. Scale bars are 10 µm.

**Fig.7.**
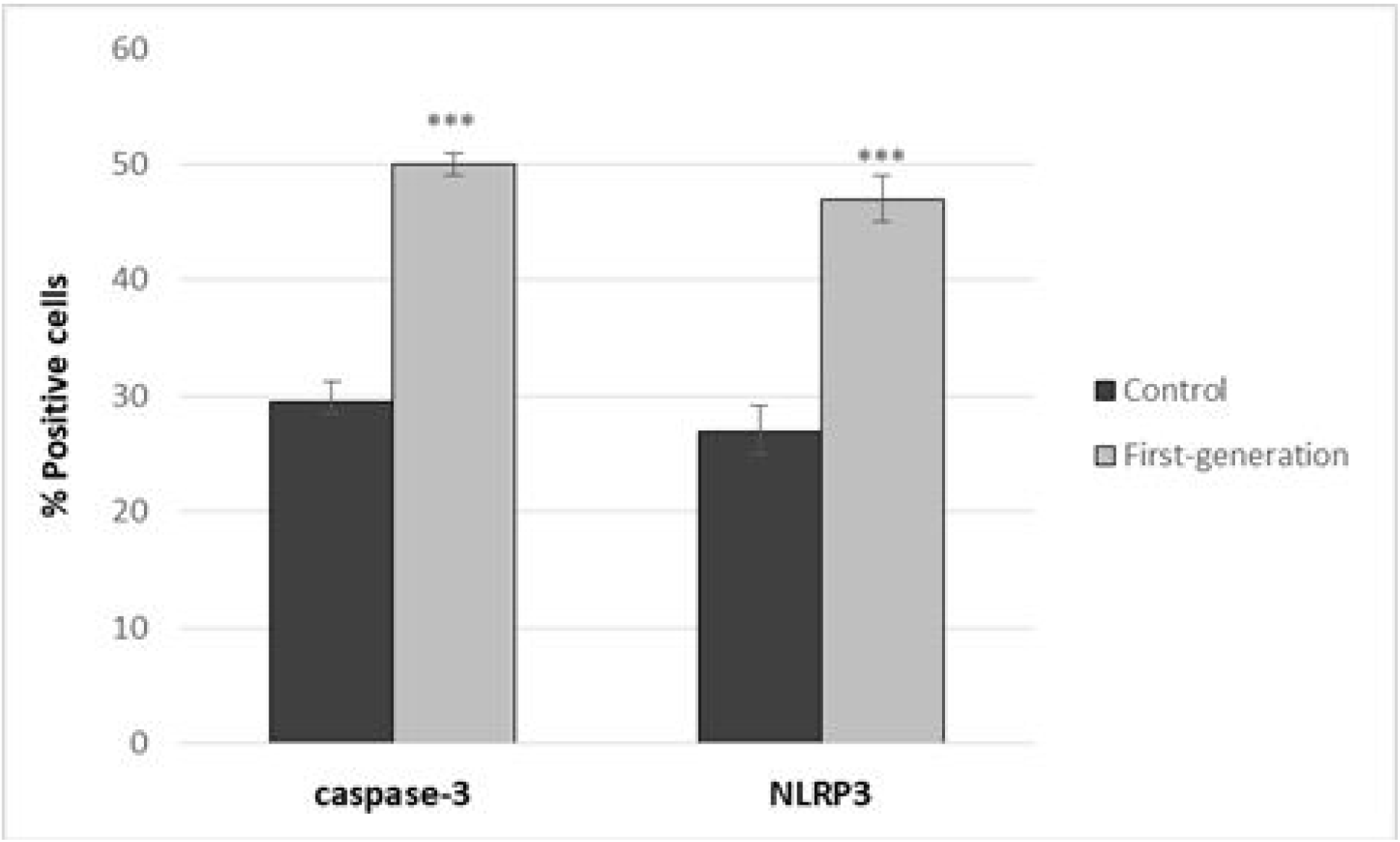
Comparison of the mean percentage of positive cells for caspase-3 and NLRP3 markers by immunocytochemical assessment. Values are reported as mean ± SD. ***p < 0.001 compared with the control group.

## Discussion

Our findings showed that maternal separation stress experienced by parents led to significant histological alterations in the ovarian tissue of first generation, including decreased percentage of primordial follicles and increased percentage of primary follicles. Furthermore, this chronic stress increased ROS production and reduced concentrations of ATP and GPx in first generation. Also, expression of cytokines and genes involved in inflammation and apoptosis including NLRP3, caspase-1, TLR4, TNFα, IL-1β, IL-18 and BCL2 were significantly affected in the first generation group. Our results also showed that maternal separation stress experienced by parents significantly increased percentage of caspase-3 and NLRP3 positive cells in the ovarian tissue of first generation.

Our previous study showed that maternal separation stress significantly affected the components of the NLRP3 inflammasome and inflammatory molecules including NLRP3, ASC, caspase-1, TLR4, IL-1β, IL-18, TNFα, and caspase-3 in the stressed mice. Furthermore, this chronic stress affected mitochondrial activation, ROS production and ATP level. Also, maternal separation stress can affect the levels of mRNA expression of BAX and BCL2 genes which play key roles in apoptosis pathways. Here, the observations that maternal separation stress experienced by parents induces alterations in ovarian tissue of first generation are novel and have not been previously reported.

According to previous studies, some parents’ experiences may not only affect the phenotype of parents but also alter the reaction to environmental impacts in the offspring (14). Franklin et al. reported that the adverse effects of chronic stress during early postnatal life can be transmitted to the following generations. They stated that transgenerational transmission of stress-induced behavioral changes in offspring can take place in a sex-dependent manner and through males. In addition, they reported that the possible mechanism for transgenerational transmission of stress-induced behavioral changes is DNA methylation alterations in the stressed male germline that are detected in the male germline and brain of following generation (22). It is well known that environmental influences including manipulation of hormones, nutritional and chemicals factors can affect mammalian DNA methylation. In addition, it can be associated with various disorders such as psychiatric, metabolic and immune disorders, as well as brain diseases and cancer (23-26).

Our findings showed that maternal separation stress experienced by parents led to histological changes in the ovarian tissue, including reduced percentage of primordial follicles and increased percentage of primary follicles. In addition, percentage of secondary and graafian follicles increased in the first-generation group, although this difference was not significant. Stress likely can accelerate the developmental process of primordial follicles to subsequent follicles. It appears that maternal separation stress experienced by parents may decrease the ovarian reserve and shorten the female reproductive lifespan.

In present study, maternal separation stress experienced by parents resulted in increase of ROS production and a reduction in GPx concentration. It is well known that ROS have certain physiological roles in the female reproduction process such as folliculogenesis, maturation of oocyte, fertilization and development of embryo. However, increase of ROS production and oxidative stress can influence the capacity of fertilization and the female reproductive lifespan (27). Moreover, increased production of ROS and oxidative stress can result in mitochondrial dysfunction, apoptosis and inflammation in the ovary. In this regard, our results also confirmed that the expression of cytokines and genes involved in the inflammation and apoptosis were affected in the first generation group.

According to the role of mitochondria in activation of NLRP3 inflammasome, damage to mitochondria can activate apoptosis and NLRP3 inflammasome. It is evidenced that mitochondrial ROS and release of oxidized mitochondrial DNA into cytosol can trigger activation of NLRP3 inflammasome (28-31). Furthermore, it has been suggested that BCL2 and TNF that play roles in apoptosis and inflammation respectively, can regulate the NLRP3 inflammasome activation (28, 32-34). Given these data, our results, plus the well-known effects of stress on the neuroendocrine system, it seems that maternal separation stress experienced by parents can affect the components of NLRP3 inflammasome and molecules involved in inflammation with change in ROS, BCL2 and TNFα levels and subsequently result in increased production of IL-1β and IL-18. Although inflammatory process is necessary for reproductive processes such as ovulation, menstruation and implantation, unbalanced inflammatory reaction can disturb the normal ovarian function (35, 36). On the other hand, it has been described that IL-1 can increase the expression of inflammatory genes and induce the apoptotic pathways that can result in the depletion of ovarian reserve (37). Also, Hussein et al. reported that mitochondrial pathway or binding of death receptors to TNF-α and Fas ligand can lead to follicular atresia (38). Given to these data and our findings, it can be concluded that maternal separation stress experienced by parents through inflammatory and apoptosis activation may adversely affect the process of folliculogenesis and follicle numbers.

Bousalham et al. showed that maternal separation stress led to depression and anxiety as well as reduction of the litter size on mother who is exposed to separation stress. Furthermore, maternal separation stress can affect subsequent generations (3). It seems that there is a relationship between the reproductive system and the pathophysiology of mood disorders. The manipulation of early environment can affect the developmental programming of the HPA axis. The timing and severity of the early manipulation as well as the sex of fetus or neonate influence the phenotype of HPA axis function (39). Clinical studies have confirmed that reduction in the hormone estrogen, an essential hormone for regulation of HPA axis, leads to increase of depression rates (40). It is stated that when the level of estrogen reduced, dysfunction of HPA axis and reproductive problems lead to mood disorders and reduction of the litter size (3).

According to our studies, early postnatal stress can induce alterations in ovarian tissue not only in the stressed mice but also in their first generation but its mechanisms are not well understood. Furthermore, these results support the hypothesis that individual risk for ovarian disorders may be affected not only on one’s own experiences but also on the parents’ experiences. Understanding the mechanisms by which maternal separation stress experienced by parents induces harmful effects on ovarian function will provide essential information for the female reproductive health of next generation.

## Conclusion

Overall, these results are the first to reveal that maternal separation stress experienced by parents can affect ovarian function in first generation. Our findings suggest that maternal separation stress experienced by parents may influence activation of inflammatory response in the ovarian tissue of their first generation which may induce apoptosis and consequently disturb folliculogenesis process. Stress-induced ovarian tissue alterations can be transmitted across first generations. Nevertheless, further studies are needed to determine the exact effects of maternal separation stress experienced by parents on ovarian function and fertility across generations.

## Acknowledgments

This research has been funded by Tehran University of Medical Sciences (TUMS); grant no. 32229. The authors thank Tehran University of Medical Sciences for its support.

